# An approach for normalization and quality control for NanoString RNA expression data

**DOI:** 10.1101/2020.04.08.032490

**Authors:** Arjun Bhattacharya, Alina M. Hamilton, Helena Furberg, Eugene Pietzak, Mark P. Purdue, Melissa A. Troester, Katherine A. Hoadley, Michael I. Love

## Abstract

The NanoString RNA counting assay for formalin-fixed paraffin embedded samples is unique in its sensitivity, technical reproducibility, and robustness for analysis of clinical and archival samples. While commercial normalization methods are provided by NanoString, they are not optimal for all settings, particularly when samples exhibit strong technical or biological variation or where housekeeping genes have variable performance across the cohort. Here, we develop and evaluate a more comprehensive normalization procedure for NanoString data with steps for quality control, selection of housekeeping targets, normalization, and iterative data visualization and biological validation. The approach was evaluated using a large cohort (N = 1,649) from the Carolina Breast Cancer Study, two cohorts of moderate sample size (N = 359 and 130), and a small published dataset (N = 12). The iterative process developed here eliminates technical variation (e.g. from different study phases or sites) more reliably than the three other methods, including NanoString’s commercial package, without diminishing biological variation, especially in long-term longitudinal multi-phase or multi-site cohorts. We also find that probe sets validated for nCounter, such as the PAM50 gene signature, are impervious to batch issues. This work emphasizes that systematic quality control, normalization, and visualization of NanoString nCounter data is an imperative component of study design that influences results in downstream analyses.

## INTRODUCTION

The NanoString nCounter platform offers a targeted strategy for gene expression quantification using a panel of up to 800 genes without requiring cDNA synthesis or amplification steps [1]. The technology offers advantages in sensitivity, technical reproducibility, and strong robustness for profiling formalin-fixed, paraffin-embedded (FFPE) samples [2]. Given these advantages, nCounter is increasingly used for longitudinal studies involving FFPE samples carried out over several years [3] and diagnostic assays in clinical settings [4,5].

Proper normalization and quality control of gene expression is necessary prior to statistical analysis to reduce unwanted variation that may be associated with technical batches or RNA degradation from sample fixation [6,7]. While some sources of variation can be enumerated a priori (e.g. different research centers, batches over time, or RNA preservation methods), not all can be captured. In all cases, it is advisable to define a quality control and normalization pipeline to detect and account for technical variation in downstream statistical modeling. All normalization methods deal with a trade-off between bias that needs correction and bias or variance that may be introduced in normalization [8].

Many approaches have been developed to normalize nCounter data. NanoString provides two forms of normalization in its commonly-used nSolver Analysis Software [9]: (A) a graphical user interface with optional background correction and positive-control and housekeeping gene normalization and (B) the Advanced Analysis tool, which draws on the NormqPCR R package [10,11] to select co-expressed housekeeping genes prior to normalization. The NanoStringNorm package implements the nSolver algorithms in R [12]. The R packages NanoStringDiff and RCRnorm use hierarchical modeling methods that incorporate information from the positive, negative, and housekeeping controls for normalization [13,14]. The NACHO R package proposes a simple quality control and visualization pipeline that precedes normalization using either NanoStringNorm or NanostringDiff [15], though, without post-normalization visualization to assess normalization quality. When technical replicates are available, a method from Molania et al, Remove Unwanted Variation-III (RUV-III), can be used along with an iterative normalization process where several parameters (i.e. number of housekeeping genes, number of detected outliers, number of dimensions of technical noise) are tuned with relevant visual and biological checks [7]. RUV-III normalization frequently outperformed nSolver normalization by more efficiently removing technical sources of variation while preserving biological variation [7]. Since many cohorts do not have technical replicates, we extend Molania et al’s iterative framework using RUVSeq [6–8], a precursor of RUV-III.

Here, we provide a framework for the quality control and normalization of mRNA expression count data from the NanoString nCounter platform, using a large dataset (N = 1,649) of breast tumor expression from the Carolina Breast Cancer Study (CBCS) and three other cohorts of differing sample size (N = 12, 130, and 359). We illustrate some of the pitfalls in the nSolver method of background correction and positive control normalization, provide an alternative approach that uses RUVSeq [6,8], and benchmark our framework against other normalization methods [9,13,14]. We find that, especially in longitudinal, multi-phase or multi-site cohorts, RUVSeq outperforms nSolver in removing differences across technical sources of variation. Lastly, we provide quality checks for normalization and outline the impact of proper normalization on inference for biological associations and expression-based disease subtyping.

## MATERIAL AND METHODS

### Data collection

We used four cohorts with nCounter gene expression data to evaluate differences between normalization procedures. Cohort details and the normalization parameters for each cohort are given below and summarized in **Supplemental Table S1**.

#### CBCS gene expression data

The Carolina Breast Cancer Study (CBCS) is a multi-phase cohort of women with breast cancer in North Carolina. Samples were collected during three study phases: Phase 1 (1993-1996), Phase 2 (1996-2001), and Phase 3 (2008-2013). Paraffin-embedded tumor blocks were reviewed and assayed for gene expression using the NanoString nCounter system as discussed previously [3,16,17]. Study phase gives the relative age of the tumor block. In total, 1,649 samples from patients with invasive breast cancer from CBCS, across all three study phases, were analyzed on a custom panel of 417 genes. All assays were performed in the Translational Genomics Laboratory (TGL) at the University of North Carolina at Chapel Hill (UNC). After quality control and normalization, 1,264 samples remained in the nSolver-normalized data, and 1,219 samples remained in the RUVSeq-normalized data. This dataset was used to benchmark against NanoStringDiff [13] and RCRnorm [14], using the same 1,264 samples in the nSolver-normalized set.

#### Bladder tumor gene expression data

FFPE Biospecimens from 42 samples of NMIBC from UNC (Chapel Hill, NC) and 88 samples from a study conducted by the Memorial Sloan Kettering Cancer Center (New York, NY) with non-muscle invasive bladder cancer (NMIBC) were analyzed. RNA was isolated using the RNeasy FFPE Kit (Qiagen) at UNC and NanoString assays were performed at the TGL at UNC using a custom codeset consisting of 440 endogenous and 6 housekeeping genes. After quality control and normalization, 86 samples remained in both the nSolver-normalized and RUVSeq-normalized datasets.

#### Kidney tumor gene expression data

This study includes 359 samples from patients with clear cell renal cell carcinoma (CCRCC) with fresh-frozen tissue collected as part of a large case-control study of kidney cancer conducted in central and eastern Europe [18]. Slides for each case were reviewed by a pathologist to assess tumor stage and grade [19]. Manual microdissection was performed to remove non-tumor tissue. Frozen sections were placed directly in Trizol reagent (Invitrogen, Carlsbad, CA), homogenized for 2 minutes on ice, and RNA was isolated using the manufacturer’s protocol. NanoString assays were performed at UNC TGL using a custom codeset consisting of 62 endogenous and 6 housekeeping genes commonly studied in kidney cancer. After quality control and normalization, 331 samples remained in both the nSolver- and RUVSeq-normalized data.

#### Sabry et al gene expression data

We downloaded raw RCC files from Sabry et al [20] from the NCBI Gene Expression Omnibus (GEO) with accession number GSE130286 and imported them using functions in NanoStringQCPro [21]. This dataset comprised of 12 samples, all of which remained after normalization with both procedures. The dataset measured 706 endogenous genes with 40 housekeeping genes from the NanoString nCounter Human Myeloid Innate Immunity Panel [20].

### Quality control and normalization

The full quality control and normalization process using nSolver and RUVSeq is summarized in **Figure 1**, starting with familiarization of the raw data (**Figure 1.1)**, technical quality control (**Figure 1.2**), pre-normalization assessment of housekeeping genes (**Figure 1.3**) and data visualization to detect problematic samples and assess whether flagged samples should be removed (**Figure 1.4**). Normalization is performed with either nSolver or RUVSeq (**Figure 1.5**), and the processed expression data is assessed for validity through relevant visualization and biological checks (**Figure 1.6**). If validation is unsatisfactory and technical variation is still present, this process is iterated.

**Figure 1:**
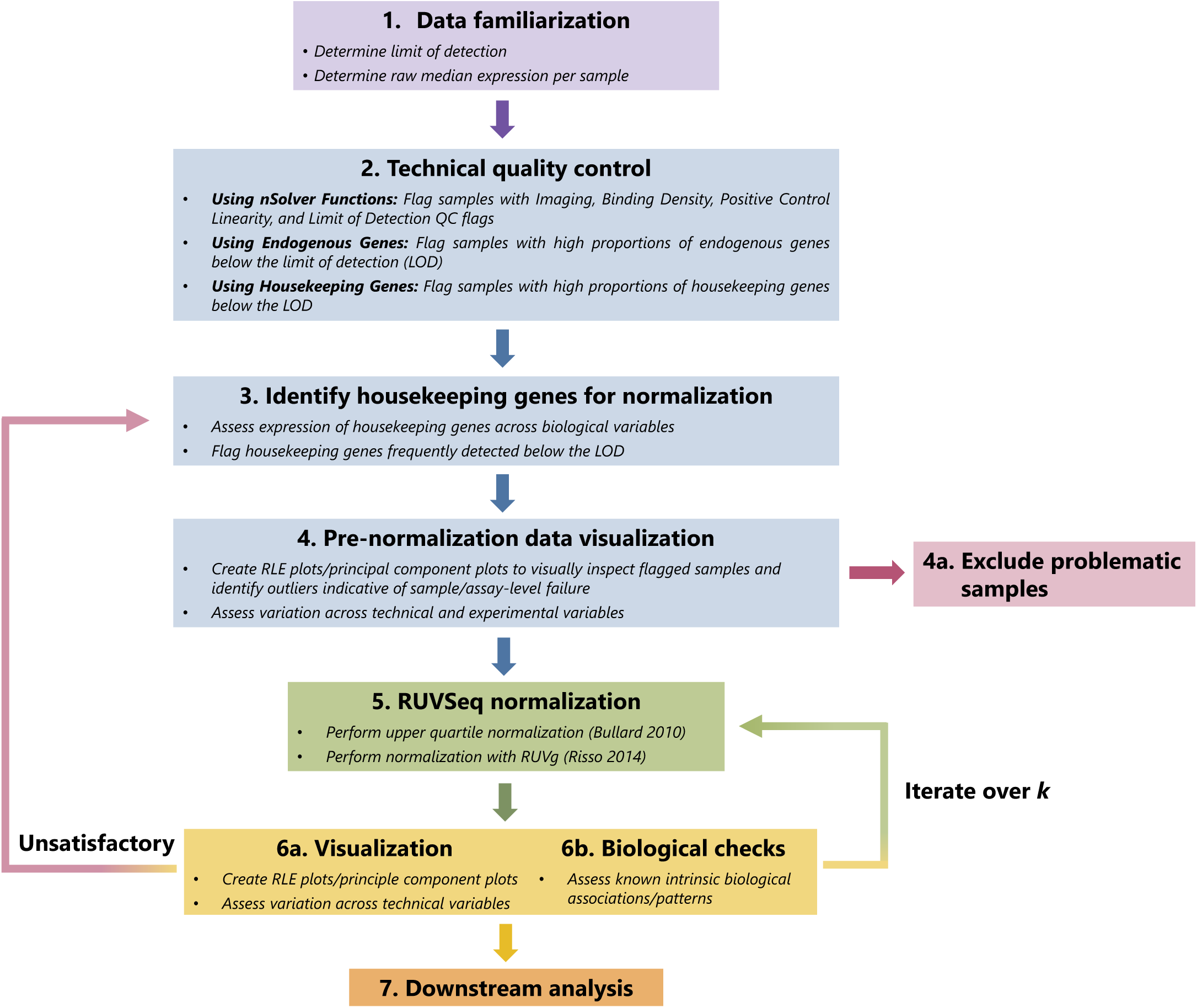
Graphical summary of RUVSeq normalization pipeline. The quality control and normalization process starts with familiarization with the data (**Step 1**) and technical quality control to flag samples with potentially poor quality (**Step 2**). After a set of housekeeping genes are selected (**Step 3**), important unwanted technical variables are also investigated through visualization techniques (**Step 4**). Problematic samples (e.g. those that are flagged multiple times in technical quality control checks) are excluded. Next, the data is normalized using upper quartile normalization and RUVSeq (**Step 5**), and the normalized data is visualized to assess the removal of unwanted technical variation and retention of important biological variation (**Step 6**). Steps 3—6 are iterated until technical variation is satisfactorily removed, changing the set of housekeeping genes or the number of dimensions of unwanted technical variation (*k*) estimated using RUVSeq. This data can then be used for downstream analysis (**Step 7**).

#### Technical quality controls flags

nSolver provides quality control (QC) flags to assess the quality of the data for imaging, binding density, linearity of the positive controls, and limit of detection. The definition and implementation of this QC is summarized in nSolver [9] and NanoStringNorm [12] documentation. We mark any sample that is flagged in at least one of these four QC assessments as technical quality control. We use these QC flags in both nSolver normalization and RUVSeq normalization.

#### Below limit of detection quality control

We use high proportions of both endogenous and housekeeping genes below the limit of detection (LOD) as a QC flag to assess reduced assay or sample quality. The per-sample LOD is defined as the mean of the counts of negative control probes for a given sample. We assessed the percent of counts below the LOD in the housekeeping genes per sample to flag both poor quality samples and housekeeping genes with problems in their measurement. We used samples with all housekeeping genes above the LOD as a reference group to determine the regular distribution of genes below the LOD. Samples were flagged if they had more than one housekeeping gene below the LOD and (2) the percent of endogenous genes below the LOD was greater than the top quartile of the distribution of percent below LOD in the reference group.

#### Housekeeping gene assessment

Housekeeping genes serve two purposes: 1) for QC purposes to remove samples with overall poor quality and 2) for assessing the amount of technical variation present in the normalization procedure. NanoString documentation suggests that ideal housekeeping genes are highly expressed, have similar coefficients of variation, and have expression values that correlate well with other housekeeping genes across all samples [9,12]. Because of these definitions, these targets will ideally vary only due to the level of technical variation present. RUVSeq relies on housekeeping genes, i.e. genes not influenced by the condition of interest (e.g. cancer subtype), with no assumptions on co-expression of all housekeeping genes. To assess the potential for housekeeping correction to introduce bias, housekeeping genes were assessed for differential expression across a primary biological covariate of interest (estrogen receptor status in CBCS, tumor stage in the kidney and bladder cancer data, and treatment groups in Sabry et al [20]) using negative binomial regression on the raw counts from the MASS package [22].

### nSolver normalization

#### Background correction

NanoString guidelines suggest background correction [9,12] by either subtraction or thresholding for an estimated background noise level for experiments in which low expressing targets are common, or when the presence of a transcript has an important research implication [7,12]. Data from all four cohorts considered do not necessarily fall under this criterion, and accordingly, we did not background correct by either method. To demonstrate the effect of background correction, we tested nSolver-normalized gene expression with and without background thresholding in CBCS using relative log expression (RLE) plots.

#### Positive control and housekeeping gene-based normalization

The arithmetic mean of the geometric means of the positive controls for each lane was computed and then divided by the geometric mean of each lane to generate a lane-specific positive control normalization factor [9,12]. The counts for every gene were multiplied by their lane-specific normalization factor. To account for any noise introduced into the nCounter assay by positive normalization, the housekeeping genes were used similarly as the positive control genes to compute housekeeping normalization factors to scale the expression values [9,12]. NanoString flagged samples with large housekeeping gene scaling factors (we call this a housekeeping QC flag) and large positive control scaling factors (positive QC flag) but note that samples with these flags simply indicate that a sample is divergent from other samples in the dataset and do not necessarily require removal. Pre-normalization visualization (**Figure 1.4)** is important for confirming the inclusion or removal of these samples.

### RUVSeq normalization pipeline

#### Normalization

The RUVSeq-based normalization process (**Figure 1.5**), an alternative approach to nSolver normalization, proceeds following quality control and housekeeping assessment. Distributional differences were scaled between lanes using upper-quartile normalization [23]. Unwanted technical factors were estimated in the resulting gene expression data with the RUVg function from RUVSeq [8]. Unwanted variation was estimated using the final set of endogenous housekeeping genes on the NanoString gene expression panel [24,25]. In general, the number of dimensions of unwanted variation to remove was chosen by iteratively normalizing the data for a given number of dimensions and checking for the removal of known technical factors already identified in the raw expression data (e.g. study phase), and presence of key biological variation (e.g. bimodality of ESR1 expression in the CBCS breast cancer data where estrogen receptor status is a known predominant feature). Further details about choosing this dimension are given by Gagnon-Bartsch et al and Risso et al [6,8]. DESeq2 was used to compute a variance stabilizing transformation of the original count data [25], and estimated unwanted variation was removed using the removeBatchEffects function from limma [26]. Ultimately, we removed 1, 1, 3, and 1 dimensions of unwanted variation from CBCS, kidney cancer, bladder cancer, and the Sabry et al datasets, respectively. RLE plots, principal component analysis and heatmaps were used to detect any potential outliers before and after normalization.

### Alternative normalization methods for benchmarking

Using CBCS data, we compared the normalized datasets from nSolver, RUVSeq, NanoStringDiff [13], and RCRnorm [14] with the raw data through visualization methods outlined above (**Figure 1.1 to 1.4**, RLE plots and scatter plots of principal components over important technical and biological sources of variation). Details about these methods are provided in **Supplemental Table S2**.

### Downstream analyses

We used several data visualization or benchmarking methods for each cohort.

#### Silhouette width analysis in CBCS

Silhouette width, a measure used to assess how similar a sample is to its own group (i.e. study phase) as compared to other groups, was used to determine the impact of the two normalization procedures on technical and biological variation [27]. Many samples with large silhouettes can be interpreted as indicating that the different study phases are distinct and that a batch effect is still present in the data.

#### eQTL analysis in CBCS

We assessed the additive relationship between the gene expression values and germline genotypes with linear regression analysis using MatrixEQTL [28], applying the same linear model as detailed in previous work [29]. Briefly, for each gene and SNP in our data, we constructed a simple linear regression, where the dependent variable is the scaled expression of the gene with zero mean and unit variance, the predictor of interest is the dosage of the alternative allele of the SNP, and the adjusting covariates are the top five principal components of the genotype matrix. We considered both cis- (SNP is less than 0.5 Mb from the gene) and trans-eQTLs in our analysis. We adjusted for multiple testing via the Benjamini-Hochberg procedure [30].

#### PAM50 subtyping in CBCS

We classified each subject into PAM50 subtypes using the procedure summarized by Parker et al [31,32]. Briefly, for each sample, we computed the Euclidean distance of the log-scale expression values for the 50 PAM50 genes to the PAM50 centroids for each of the molecular subtypes. Each sample was classified to the subtype with the minimal distance [31]. The PAM50 genes were clustered hierarchically for both samples and genes and visualized in heatmaps. Subtype concordance was assessed between normalization methods excluding normal-like cases.

#### RNA-seq normalization and distance correlation analysis in CBCS

We obtained a separate set of samples (not included in the analysis described above) from CBCS with both RNA-seq and nCounter expression (on a different codeset of 166 genes). We followed a standard RNA-seq normalization process with DESeq2 [25], using the median of ratios method to estimate scaling factors [24]. We calculated the distance correlation and conducted a multivariate permutation test of independence between the RNA-seq data set (subset to the overlapping genes on the NanoString codeset) with each of the nSolver-normalized and RUVSeq-normalized nCounter data using the energy package [33]. The distance correlation and associated permutation test allow for detection of non-independence across multivariate datasets of different distribution.

#### Differential expression analysis with Sabry et al. dataset [20]

We conducted differential expression analysis to compare both normalization methods in the Sabry et al. dataset [20] using DESeq2 [25], and adjusting for multiple testing with the Benjamini-Hochberg [30] procedure. We compared differential expression across IL-2–primed NK cells vs. NK cells alone and CTV-1-primed NK cells for 6 hours vs. NK cells alone.

## RESULTS

We evaluated the ability of normalization methods to remove technical variation while retaining biologically meaningful variation across four cohorts of differing sample size and varying sources of technical bias (**Supplemental Table S1**). Known sources of technical variation included age of sample (study phase) and different study sites. The cohorts varied in preservation methods; two cohorts used fresh-frozen specimens, while two used archival FFPE specimens. The number of genes measured for both endogenous genes and housekeeping genes also varied by study. In addition, some studies used validated and optimized code sets for specific gene signatures versus a more general code set.

In cohorts with large technical biases, RUVSeq provided superior normalization with more robust removal of technical variation and provided stronger biological associations compared to other normalization methods. In two of the datasets, we found that downstream analyses performed on data normalized with nSolver and RUVSeq detected substantially different biological associations. However, when few strong technical biases were present or if a validated and optimized code set (e.g. PAM50 genes) was used, nSolver and RUVSeq performed comparably.

### Case study: Carolina Breast Cancer Study

#### Evaluation of background correction

Background thresholding led to increased per-sample variance while per-sample medians remained relatively similar (**Supplemental Figure S1A**). The distributions of per-sample median expression values were more right-skewed (greater mean than median) when using background thresholding prior to normalization compared to not using background thresholding (**Supplemental Figure S1B**). Based on this analysis, we did not perform background correction prior to normalization for all cohorts analyzed.

#### Quality assessment of expression levels using LOD of housekeeping genes

We used the housekeeping genes to assess if the lack of expression of endogenous genes was due to biology or due to technical failures. We compared the level of missing endogenous genes in samples with all housekeeping genes present to those with increasing number of housekeeping genes below LOD. There was a strong positive correlation for increasing proportions of genes below the LOD in both the endogenous and housekeeping genes (**Figure 2A**;**Supplemental Figure S2)**. Samples with higher numbers of genes below the LOD were from earlier phases of CBCS (i.e. Phase 1 from 1993-1996 and Phase 2 from 1996-2001), and thus associated with sample age (**Figure 2A**;**Supplemental Figure S3**). Samples with a higher proportion of endogenous genes below the LOD had increased numbers of QC flags as well (**Supplemental Figure S2**).

**Figure 2:**
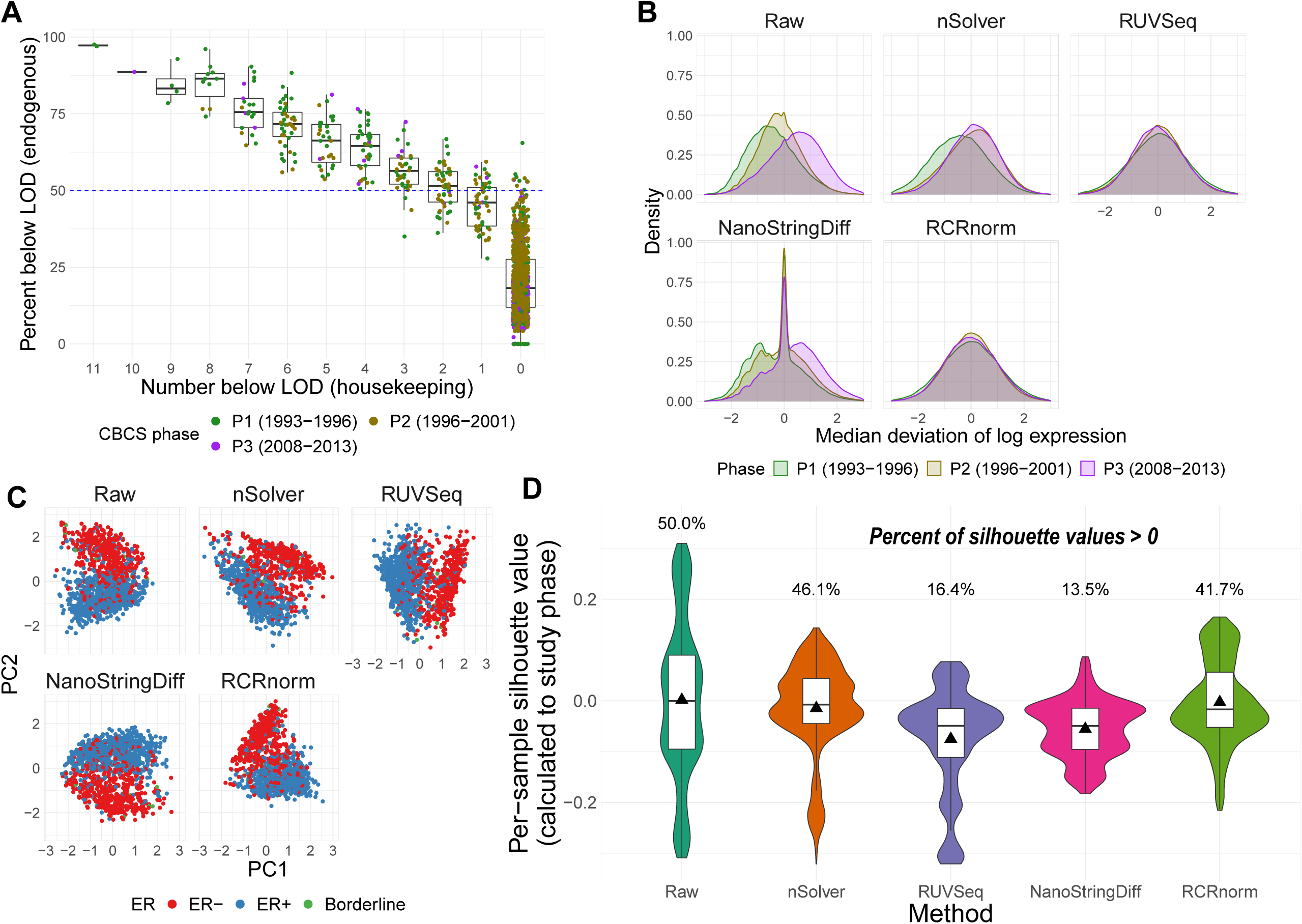
Quality control and normalization validation in CBCS. **(A)** Boxplot of percent of endogenous genes below the limit of detection (LOD) (*Y*-axis) over varying numbers of the 11 housekeeping genes below LOD (*X*-axis), colored by CBCS study phase. Note that the *X*-axis scale is decreasing. **(B)** Kernel density plots of deviations from median per-sample log_2_-expression from the raw, nSolver-, RUVSeq-, NanoStringDiff-, and RCRnorm-normalized expression matrices, colored by CBCS study phase. **(C)** Plots of the first principal component (*X*-axis) vs. second principal component (*Y*-axis) colored by estrogen receptor subtype of the raw, nSolver-, RUVSeq-, NanoStringDiff-, and RCRnorm-normalized expression data. **(D)** Violin plots of the distribution of per-sample silhouette values, as calculated to study phase, using raw, nSolver-, RUVSeq-, NanoStringDiff-, and RCRnorm-normalized expression. The boxplot shows the 25% quartile, median, and 75% quartile of the distribution, and the plotted triangle shows the mean of the distribution.

#### Evaluation of normalization methods

We benchmarked RUVSeq and nSolver with two other normalization methods, NanoStringDiff [13] and RCRnorm [14]. We observed differences across the four normalization strategies (described in **Supplemental Table S2**), namely greater remaining technical variation using nSolver and NanoStringDiff than RCRnorm and RUVSeq **(Figure 2B-D)**. A large portion of the variation in the raw expression could be attributed to study phase **(Supplemental Figure S4A).** While all methods reduced study phase associated variation compared to the raw data, there were considerable differences in the deviations from the median log-expressions in the nSolver- and NanoStringDiff-normalized expression that are not present in the RUVSeq- and RCRnorm-normalized data **(Figure 2B)**. The nSolver and NanoStringDiff methods retained technical variation, either not fully corrected or re-introduced during the nSolver normalization process.

We examined the ability of each normalization method to retain biological variation. Estrogen Receptor (ER) status is one of the most important clinical and biological features in breast cancer and is used for determining course of treatment [34,35]. ER status drives many of the molecular classifications [36–38] and even drives separate classification of breast tumors in TCGA’s pan-cancer analysis of 10,000 tumors [39]. In the raw expression, variation due to ER status was captured in PC2 rather than PC1 (study age); however, after RUVSeq-normalization, ER status was reflected predominantly in PC1 (**Figure 2C**). In the nSolver-, NanoStringDiff-, and RCRnorm-normalized data, ER status was shared between PC1 and PC2, suggesting that unresolved technical variation was still present. RUVSeq demonstrated effective removal of technical variation and boosting of the true biological signal. The PAM50 molecular subtypes [31], which are also linked with ER status, were also clearly separated by PC1 for RUVSeq-normalized data, but this was not thess case for nSolver-, NanoStringDiff-, or RCRnorm-normalization (**Supplemental Figure S4B**). These results suggest that RUVSeq-normalization best balances the removal of technical variation with the retention of important axes of biological variation, with RCRnorm showing better performance than nSolver and NanoStringDiff, but not superior to RUVSeq. A significant disadvantage of RCRnorm is its computational cost: RCRnorm was unable to run on the CBCS dataset (N = 1278 after QC) on a 64-bit operating system with 8 GB of installed RAM, requiring RCRnorm-normalization to be performed on a high-performance cluster. We summarize the maximum memory used by method in CBCS in **Supplemental Table S2**.

We used silhouette width to assess extent of unwanted technical variation from study phase remaining by the normalization methods. Larger positive silhouette values indicate within-group similarity (i.e. samples clustering by study phase). Per-sample silhouettes across the alternatively normalized datasets showed that RUVSeq best addressed the largest source of technical variation identified in the raw data (**Figure 2D**; **Supplemental Figure S5A**) while also not removing a significant portion of biological variation (**Supplemental Figure S5B**). NanoStringDiff also demonstrated less similarity of samples across study phase similar to RUVSeq but removed biologically relevant similarity of samples grouped by ER status. Due to the performance of NanoStringDiff and computational limitations of RCRnorm, for subsequent analyses and datasets, we only illustrate differences between nSolver- and RUVSeq-normalized data.

#### Genomic analyses and expression profiles across normalization methods

We evaluated the impact of normalization choice on downstream analyses including eQTLs, PAM50 molecular subtyping, known expression patterns, and similarity to RNA-seq data. In a full cis-trans eQTL analysis accounting for race and genetic-based ancestry, we found considerably more eQTLs using nSolver as opposed to RUVSeq, thresholding at nominal *P* < 10^−3^ (2,050 vs. 1,143). We identified strong cis-eQTL signals in both normalized datasets; however, stronger FDR values were identified with RUVSeq (**Figure 3A**, densely populated around the 45-degree line). We observed considerably more trans-eQTLs using nSolver, including a higher proportion of trans-eQTLs across various FDR-adjusted significance levels (**Figure 3B; Supplemental Figures S6-S7**). We suspected that spurious trans-eQTLs may have resulted from residual technical variation in expression data that was confounded with study phase, subsequently being identified as a QTL due to ancestry differences across study phase. In cross-chromosomal trans-eQTL analysis, distributions of absolute differences in minor allele frequency (MAF) for trans-eSNPs across women of African and European ancestry were wide for both methods (**Supplemental Figure S7)**. However, we observed substantially more trans-eSNPs with moderate absolute MAF differences across study phase with nSolver, compared to RUVSeq. This provides some evidence for the presence of residual confounding technical variation in the nSolver-normalized expression data leading to spurious trans-eQTL results (with a directed acyclic graph for this hypothesis in **Supplemental Figure S8**), though we cannot confirm this with eQTL analysis alone.

**Figure 3:**
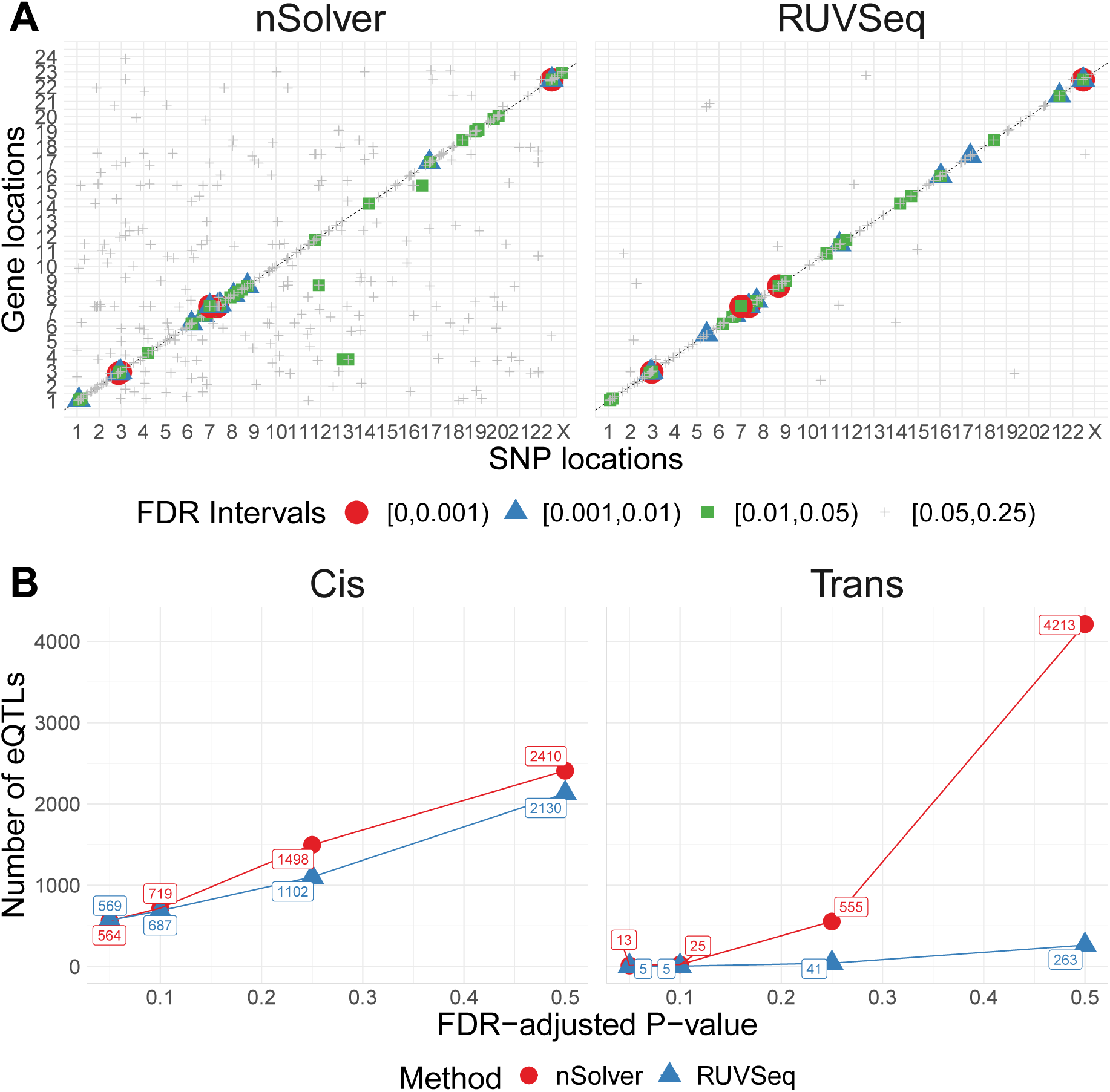
eQTL analysis in CBCS. **(A)** Cis-trans plots of eQTL results from nSolver-normalized (left) and RUVSeq-normalized data with chromosomal position of eSNP on the *X*-axis and the transcription start site of associated gene in the eQTL (eGene) on the *Y*-axis. Points for eQTLs are colored by FDR-adjusted *P*-value of the association. **(B)** The dotted line provides a 45-degree reference line for cis-eQTLs. Number of cis- (left) and trans-eQTLs (right) across various FDR-adjusted significance levels. The number of eQTLs identified in nSolver-normalized data is shown in red and the number of eQTLs identified in RUVSeq-normalized data is shown in blue.

We compared each normalization method for the ability to classify breast cancer samples into PAM50 intrinsic molecular subtype using the classification scheme outlined by Parker et al [31]. Our PAM50 subtyping calls were robust across normalization methods with 91% agreement and a Kappa of 0.87 (95% CI (0.85, 0.90)) (**Supplemental Table S3**). Among discordant calls, approximately half had low confidence values from the subtyping algorithm, and half had differences in correlations to centroids less than 0.1 between the discordant calls (data not shown). Most of these discordant calls were among HER2-enriched, luminal B and luminal A subtypes, which are molecularly similar [40].

We observed noticeable differences between the RUVSeq- and nSolver-normalized gene expression when visualized after hierarchical clustering via heatmaps, similar to the principal component analysis. Using this method, we identified 14 additional samples with strong technical errors in the nSolver-normalized data not previously marked by QC flags (**Supplemental Figure S9**), emphasizing the need for post-normalization data visualization. In early breast cancer clustering papers, the first major division was by ER status separating basal-like and HER2-enriched molecular subtypes (predominantly ER-negative) from luminal A and B molecular subtypes (predominantly ER-positive) [31]. This pattern was observed in RUVSeq-data but only partially preserved with nSolver normalization (**Supplemental Figure S9**). Rather, nSolver data clustering was driven by a combination of ER status and study phase. Study phase dominated two of the groups and were formed by Phase 1 and Phase 3 samples, respectively—samples with a 10+ year difference in age.

Lastly, we compared normalization choices for NanoString data to RNA-seq data performed on the same samples. CBCS collected RNA-seq measurements for 70 samples that have data on a different nCounter codeset (162 genes instead of 417) and RNA-seq normalized using standard procedures. A permutation-based test of independence using the distance correlation [33,41] revealed that the distance correlation between the RNA-seq and nSolver data was small and near 0 (distance correlation = 0.051, *P* = 0.24) while the distance correlation between the RNA-seq and RUVSeq-data was larger (distance correlation = 0.36, *P* = 0.02). The permutation-based test rejected the null hypothesis of independence (distance correlation of zero for unrelated datasets) between RUVSeq-normalized nCounter data and RNA-seq data but fails to reject the null hypothesis for nSolver-normalization nCounter and RNA-seq data. We conclude that RUVSeq produced normalized data with closer relation to the RNA-seq, in terms of distance correlation and test of independence, compared to nSolver.

### Case study: differential expression analysis in natural killer cells

We looked at the impact of the two normalization methods in a small cohort (*N* = 12) on DE analysis across natural killer (NK) cells primed for tumor-specific cells and cytokines from Sabry et al [20]. RLE plots before and after normalization showed minor differences between the two normalization methods (**Supplemental Figure S9**).

Using DESeq2 [25], we identified genes differentially expressed in NK cells primed by CTV-1 or IL-2 cytokines compared to unprimed NK cells at FDR-adjusted *P* < 0.05. The two normalization methods led to a different number of differentially expressed genes with a limited overlap of significant genes by both methods (**Figure 4A).** The raw *P*-value histograms from differential expression analysis using nSolver-normalized expression exhibited a slope toward 0 for *P*-values under 0.3, which can indicate issues with unaccounted-for correlations among samples [42], such as residual technical variation. The distributions of *P*-values using the RUVSeq-normalized data were closer to uniform throughout the range [0,1] for most genes (**Figure 4B**). While the log_2_-fold changes were correlated between the two normalization procedures, the genes found to be differentially expressed only with nSolver-normalized data tended to have large standard errors with RUVSeq-normalized data and therefore not statistically significant using RUVSeq (**Figure 4C**). These differences in DE results emphasize the importance of properly validating normalization prior to downstream genomic analyses.

**Figure 4:**
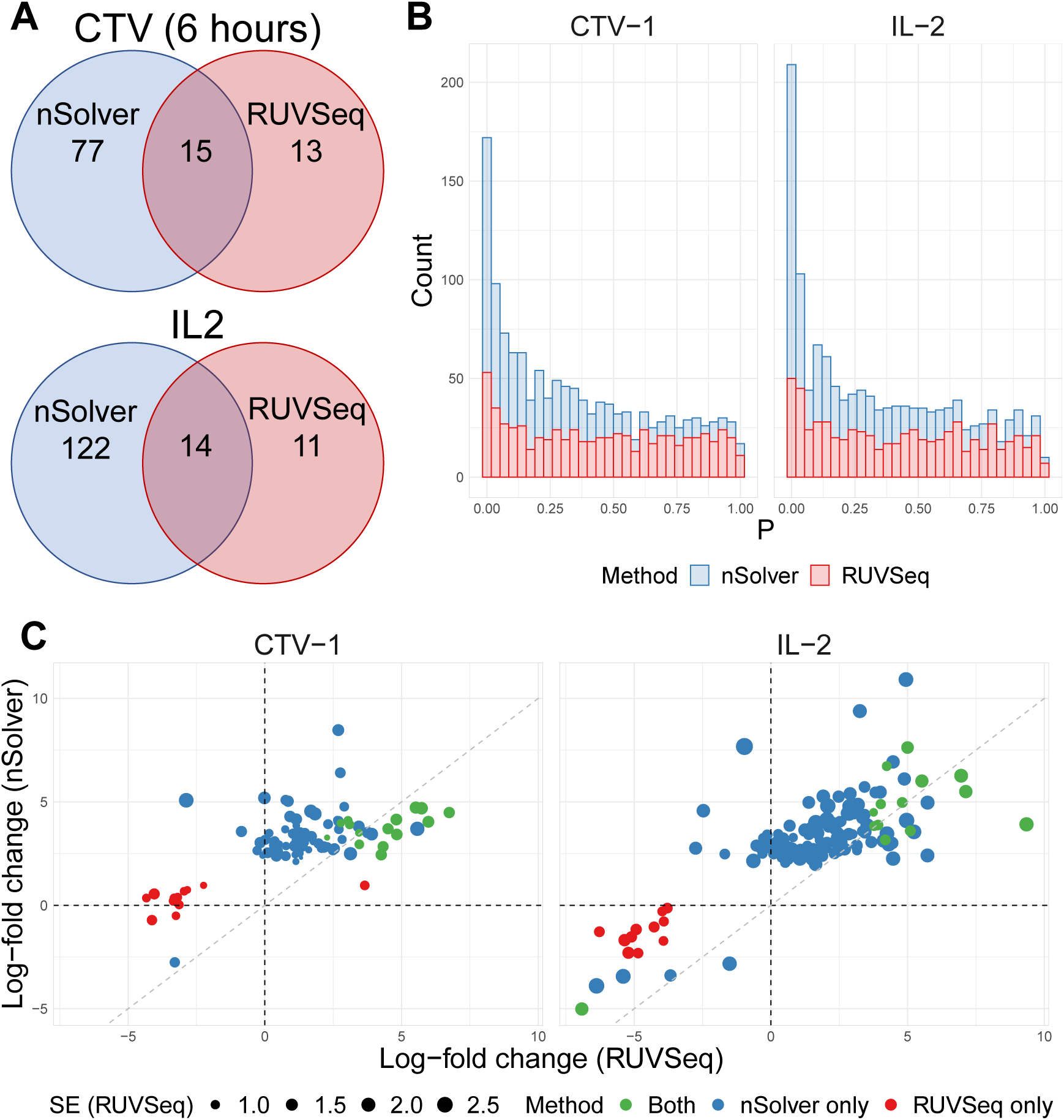
Differential expression analysis from Sabry et al [20]. **(A)** Venn diagram of the number of differentially expressed genes using nSolver-normalized (blue) and RUVSeq-normalized data (red) across comparisons for IL-2-primed (top) and CTV-1-primed NK cells (bottom). **(B)** Raw *P*-value histograms for differential expression analysis using nSolver-normalized (blue) and RUVSeq-normalized (red) data across the two comparisons. **(C)** Scatterplots of log_2_-fold changes from differential expression analysis using RUVSeq-normalized data (*X*-axis) and nSolver-normalized data (*Y*-axis) for any gene identified as differentially expressed in either one of the two datasets. Points are colored by the datasets in which that given gene was classified as differentially expressed. The size of point reflects the standard error of the effect size as estimated in the RUVSeq-normalized data. *X*= 0, *Y* = 0, and the 45-degree lines are provided for reference.

### Case study: bladder cancer gene expression

RUVSeq reduced technical variation (study site) while maintaining the biological variation (tumor grade). RUVSeq data showed the most homogeneity in per-sample median deviation of log-expressions compared to raw and nSolver data (**Figure 5A**). The first principal component of nSolver data had significant differences by study sites, which was not present in RUVSeq data (**Figure 5B**). In addition, there was a stronger biological association with tumor grade in the first principal component of expression using RUVSeq data (**Figure 5C**).

**Figure 5:**
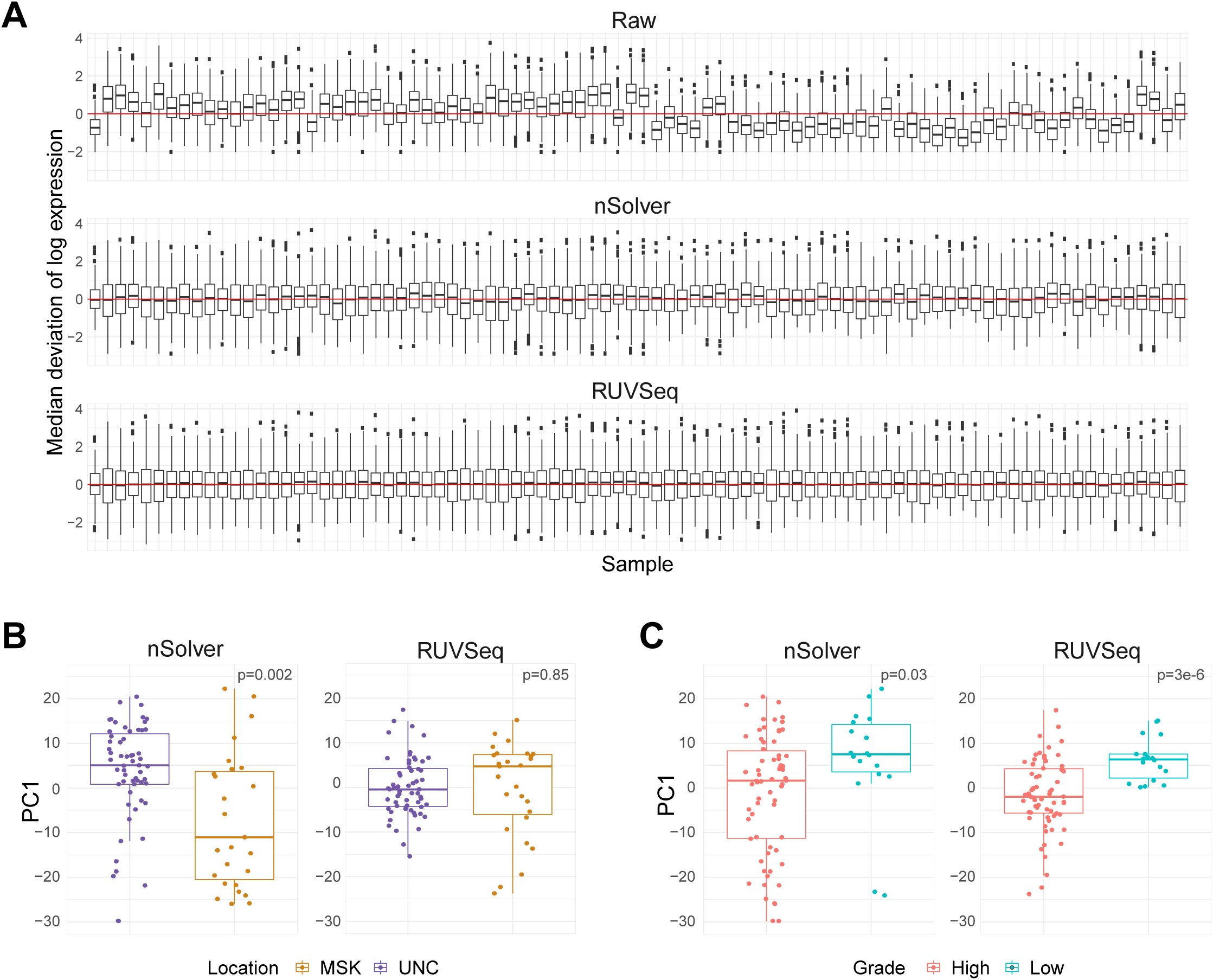
Normalization differences in bladder cancer dataset. **(A)** RLE plot from bladder cancer dataset, ordered temporally from oldest to newest sample. **(B)** Boxplot of first principal component of expression by tumor collection site (location) across nSolver- (left) and RUVSeq-normalized (right) data. Boxplot of first principal component of expression by tumor grade across nSolver- (left) and RUVSeq-normalized (right) data.

### Case study: kidney cancer gene expression

We only found subtle differences in the deviations from the median expression between the normalization procedures for the kidney cancer dataset (**Figure 6A**). This cohort did not have the same known technical variables observed in the other cohorts such as study site or sample age, and the RNA came from fresh-frozen material (**Supplemental Table S1**). We evaluated normalization methods on a source of technical variation, DV300, the proportion of RNA fragments detected at greater than 300 base pairs as a source of technical variation, and tumor stage as a biological variable of interest. The first two principal components colored by level of DV300 (**Figure 6B**) and tumor stage (**Figure 6C**) showed little difference across the two normalization methods. When there were limited sources of technical variation and a robust, high quality dataset, we found both normalization methods performed equally well.

**Figure 6:**
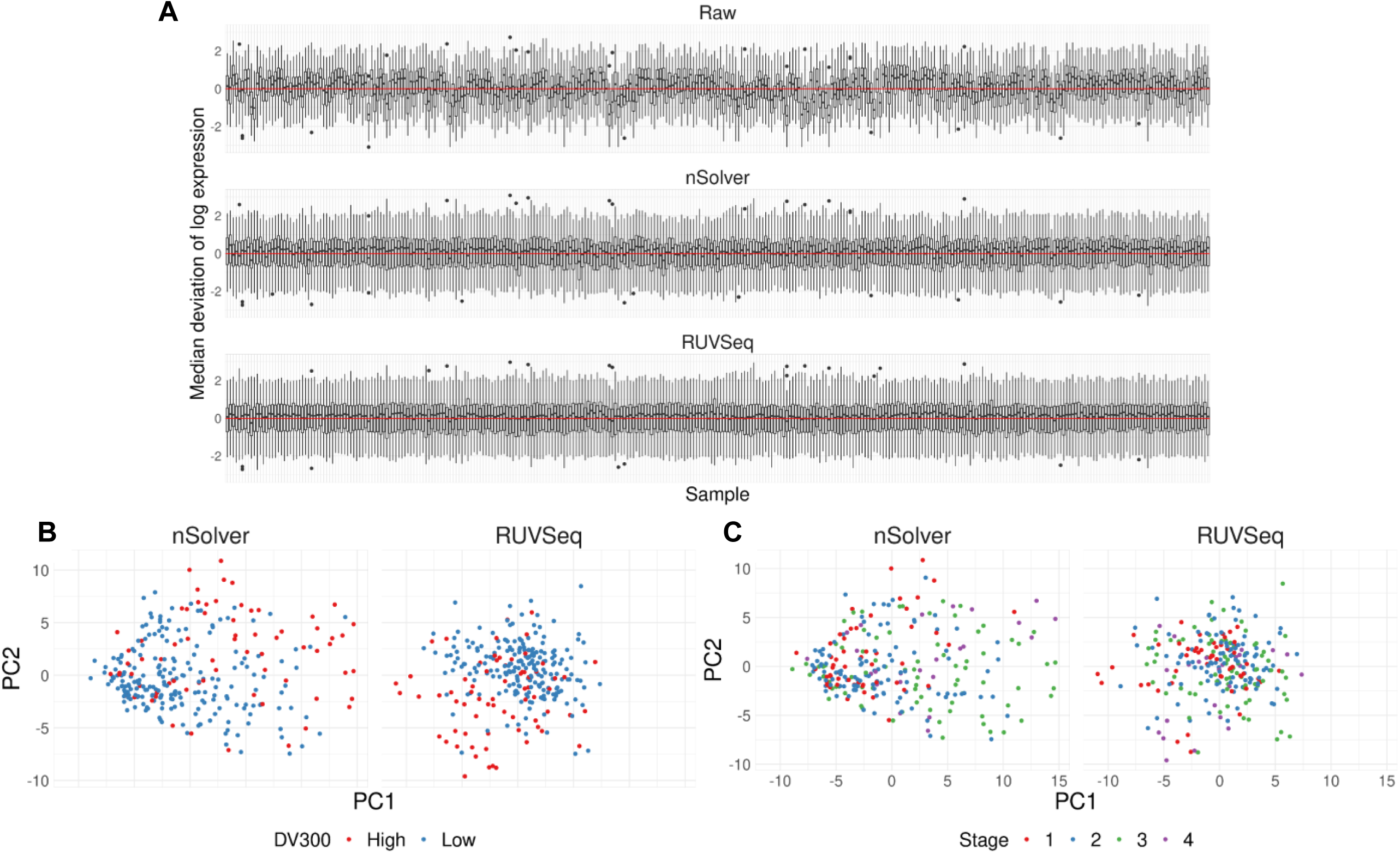
Equal performance of normalization procedures in kidney cancer dataset. **(A)** RLE plot of per-sample deviations from the median for raw, nSolver-, and RUVSeq-normalized data. **(B)** Scatter plot of the first and second principal component of nSolver- (left) and RUVSeq-normalized (right) expression, colored by high and low DV300. **(C)** Scatter plot of the first and second principal component of nSolver- (left) and RUVSeq-normalized (right) expression, colored by tumor stage.

## DISCUSSION

Proper normalization is imperative in performing correct statistical inference from complex gene expression data. Here, we outline a sequential framework for NanoString nCounter RNA expression data that provides both quality control checks, considerations for choosing housekeeping genes, and iterative normalization with biological validation using both NanoString’s nSolver software [9,12] and RUVSeq [6,8]. We show that RUVSeq provided a superior normalization to nSolver on three out of four datasets by more efficiently removing sources of technical variation, while retaining robust biological associations.We also benchmark RUVSeq-normalization with two other normalization methods implemented in R and show that RUVSeq outperformed all methods in reducing technical variation.

We observed that normalization methods were sensitive to the quality and the set of housekeeping genes. Several genes thought to behave exclusively in a “housekeeping” fashion in fact associate with biological variables under certain conditions [43] or across different tissue types [44]. A careful validation of housekeeping gene stability on a case-by-case basis and separately for new studies, considering both technical and biological sources of variation in each dataset, is therefore imperative for an optimized normalization procedure.

We developed a quality metric to assess sample quality: samples with high proportions of genes detected below the LOD in both endogenous genes and housekeepers were indicative of either low-quality samples or reduced assay efficiency. Sample age was correlated with higher proportions of genes below the LOD in both endogenous and housekeeping genes, which was likely due to RNA degradation over time. We stress that missing counts in endogenous genes alone does not suggest poor sample quality in the absence of additional QC flags but could represent genes not expressed and therefore not detected under certain biological conditions or cell types. An example includes using an immuno-oncology gene panel in a tumor sample with little to no immune cell infiltration. Conversely, many samples with counts below the LOD in both endogenous genes and housekeepers had additional quality control flags including those derived from nSolver’s assessment of data quality. We excluded these samples for analysis in both the nSolver- and RUVSeq-based procedures.

nSolver-normalized data was prone to residual unwanted technical variation when there were known technical biases, such as in CBCS and the bladder example. We checked for known biological associations that are intrinsic to the sample, as in eQTL analysis, to judge the performance of the normalization process [45,46]. A full cis-trans eQTL analysis using nSolver- and RUVSeq-normalized data showed a strong cis-eQTL signal in data from both normalization methods. We found significantly more trans-eQTLs with the nSolver-normalized data (**Figure 3**). However, many of the trans-eSNPs for the loci found with nSolver-normalized data tended to have moderate MAF differences across phase, leading us to suspect they were spurious associations driven by residual technical variation in gene expression (**Supplemental Figure 8**). Such spurious associations from population stratification have been described in many previous studies of eQTL analysis [47–50].

The choice of normalization procedure is less of a concern in cohorts with minimal sources of technical variation or in nCounter targeted gene panels that have been optimized for robust measurement across preservation methods. In the CBCS breast cancer cohort, we identified significant differences in gene expression between normalization methods across the entire gene set (417 total genes). However, PAM50 subtyping was robust across the two normalization procedures. The genes in the PAM50 classifier were selected due to their consistent measurement in both FFPE and fresh frozen breast tissues [31], suggesting that robustly measured genes may be less affected by different normalization procedures. Furthermore, we see minimal differences in residual technical variation in the kidney cancer dataset and the Sabry et al dataset, both of which were measured on either robustly validated genes or nCounter panels. The kidney cancer example had newer, fresh-frozen specimens that were profiled using a small and well-validated set of genes important in that cancer type. This dataset gives an opportunity to stress the importance of the general principles of normalization: as Gagnon-Bartsch et al and Molania et al recommend [6,7], normalization should be a part of scientific process and should be approached iteratively with visual inspection and biological validation to tune the process. One normalization procedure is not necessarily applicable to all datasets and must be re-evaluated on each dataset.

In conclusion, we outline a systematic and iterative framework for the normalization of NanoString nCounter expression data. Even without background correction, a technique which has been shown to impair normalization of microarray expression data [51,52], we believe that relying solely on positive control and housekeeping gene-based normalization may result in residual technical variation after normalization. Here, we show the merits of a comprehensive procedure that includes sample quality control checks including the addition of new checks, assessments of housekeeping genes, normalization with RUVSeq [6,8] and data analysis with popular count-based R/Bioconductor packages, as well as iterative data visualization and biological validation to assess normalization. Researchers must pay close attention to the normalization process and systematically assess pipelines that best suit each dataset.

## KEY POINTS

- The NanoString nCounter RNA counting assay, an attractive option in archived samples, has sub-optimal quality control and normalization pipelines.
- We provide an iterative framework for nCounter data with steps for quality control, normalization, and visualization/validation using RUVSeq.
- Using four real datasets, we show that our framework eliminates technical variation more reliably than other methods, including NanoString’s provided software nSolver, without diminishing biological variation.
- We stress that quality control and normalization must be emphasized in study design and evaluated using proper visualization and other checks, or else results in downstream analyses may be biased.

## AVAILABILITY

Relevant R code for these analyses are freely bundled into an R package on Github: https://github.com/bhattacharya-a-bt/NanoNormIter. R code to recreate the Sabry et al analysis and a tutorial for the iterative framework is also provided: https://github.com/bhattacharya-a-bt/CBCS_normalization/ [53]. Summary statistics for eQTL analysis are available at https://github.com/bhattacharya-a-bt/CBCS_TWAS_Paper [54], as a part of Bhattacharya et al [29].

CBCS genotype datasets analyzed in this study are not publicly available as many CBCS patients are still being followed and accordingly CBCS data is considered sensitive; the data is available from M.A.T upon reasonable request. Raw and normalized expression data from CBCS will be available on GEO upon publication. For replication or review prior to publication, this data can be accessed from GEO through a reviewer token or requested from M.A.T. Data from the bladder and kidney cancer datasets may be provided by the authors upon reasonable request.

## ACCESSION NUMBERS

Raw RCC files for nCounter expression from Sabry et al [20] are available NCBI Gene Expression Omnibus (GEO) with the accession numbers GSE130286. Raw and normalized expression data from CBCS will be available on GEO upon publication. For replication prior to publication, this data can be requested from the authors.

## SUPPLEMENTARY DATA

Document S1: Supplemental Tables and Figures

## AUTHOR BIOGRAPHICAL STATEMENT

A.B. is a Doctoral Candidate in Biostatistics, and A.M.H. is a Doctoral Candidate in Pathology and Laboratory Medicine, both at the University of North Carolina at Chapel Hill. H.F. is an Associate Attending Epidemiologist, and E.P. is a Urological Surgeon, both at Memorial Sloan Kettering Cancer Center. M.P. is a Senior Investigator in the Division of Cancer Epidemiology and Genetics, National Cancer Institute. M.A.T is a Professor of Epidemiology and Pathology and Laboratory Medicine, K.A.H. is an Assistant Professor of Genetics, and M.I.L is an Assistant Professor of Biostatistics and Genetics, all at the University of North Carolina at Chapel Hill.

## ACKNOWLEDGEMENT

We thank the Carolina Breast Cancer Study participants and volunteers. We thank Halei Benefield, Xiaohua Gao, Erin Kirk, Linnea Olsson, and Jessica Tse for their invaluable support during the research process.

