## Supplementary material for "An approach for normalization and quality control for NanoString RNA expression data": Document S1: Supplemental Tables and Figures

**Supplemental Figures**

|  | **Study/Dataset** | **Preservation Method** | **N** | **Failed QC** | **Passed QC** | **Codeset Size (genes)** | **# Available HK genes** | **# HK Genes Used** | **HK Genes Used in Normalization** | **k** | **Downstream Analysis** | **Source of Technical Variation** |
| --- | --- | --- | --- | --- | --- | --- | --- | --- | --- | --- | --- | --- |
| **RUVseq** | **Carolina Breast Cancer Study** | FFPE | 1649 | 430 | 1219 | 417 | 11 | 7 | ACTB, CLTC, GAPDH, HPRT1, PGK1, RPLP0, SF3A1 | 1 | RLE, Silhouette width, eQTL, PAM50 | Sample age |
|  | **Kidney Tumor** | Frozen | 359 | 28 | 331 | 68 | 6 | 5 | HMBS, GUSB, POLR2A, PPIA, TFRC | 1 | RLE, PCA | None known |
|  | **Bladder Tumor** | FFPE | 130 | 44 | 86 | 446 | 6 | 5 | NAGA, KU70 (aka XRCC6), GUSB, RPS10, PGK1, ACTB | 3 | RLE, PCA | Study site, sample age |
|  | **Sabry *et al.* Study** | Frozen | 12 | 0 | 12 | 706 | 40 | 40 | Refer to Sabry *et al*., | 1 | Differential expression | None known |
| **nSolver** | **Carolina Breast Cancer Study** | FFPE | 1649 | 385 | 1264 | 417 | 11 | 7 | ACTB, CLTC, GAPDH, HPRT1, PGK1, RPLP0, SF3A1 | - | RLE, Silhouette width, eQTL, PAM50 | Sample age |
|  | **Kidney Tumor** | Frozen | 359 | 28 | 331 | 68 | 6 | 5 | HMBS, GUSB, POLR2A, PPIA, TFRC | - | RLE, PCA | None known |
|  | **Bladder Tumor** | FFPE | 130 | 44 | 86 | 446 | 6 | 5 | NAGA, KU70 (aka XRCC6), GUSB, RPS10, PGK1, ACTB | - | RLE, PCA | Study site, sample age |
|  | **Sabry *et al.* Study** | Frozen | 12 | 0 | 12 | 706 | 40 | 40 | Refer to Sabry *et al*., | - | Differential expression | None known |

**Table 1**: Summary of NanoString datasets normalized by RUVseq and nSolver. We provide the study goals or endpoints considered, the preservation method of the samples, the sample size of the study, the number of samples that pass and fail quality control, the number of genes in the codeset, the number of housekeeping genes available in the given condeset, the number and names of housekeeping genes we eventually used, the number $k$ of unwanted dimensions of variation removed using RUVSeq, and known sources of technical variation.

| **Software** | **Implementation** | **Method** | **Memory Used (CBCS Data)** | **Notes** |
| --- | --- | --- | --- | --- |
| nSolver [1] | GUI with R backend | Positive- and housekeeping-control scaling | 0.41 GB (using NanoStringNorm) | Can be implemented entirely in R using the NanoStringNorm package [2] |
| RUVSeq [3,4] | R package (Bioconductor) | Factor analysis on housekeeping genes or technical replicates | 3.92 GB | RUV-III has been created by the same group for NanoString data using technical replicates [5] |
| NanoStringDiff [6] | R package (Bioconductor) | Generalized linear model | 0.37 GB | Normalized data is recommended to be used only for downstream differential expression analysis using an empirical Bayes shrinkage approach |
| RCRnorm [7] | R package (CRAN) | Bayesian random-coefficient hierarchical regression | 20.93 GB | High computational cost, even in “fast” mode |

**Table 2**: Summary of normalization software compared in benchmarking. We provide the implementation of the software, a brief summary of the methods used by the software, total memory used on a submitted job on a high performance cluster with 25 GB allocated RAM, and any miscellaneous notes about the methods (i.e. alternative implementations and disadvantages of each method). The memory used is calculated from a submitted job that processed the CBCS expression data (417 genes, 1264 samples).

**Figure 1**: **(A)** Scatter plot of per-sample median and per-sample variance of CBCS expression across raw expression (left), nSolver-normalized data with background correction (middle), and without background correction (right), with samples colored by study phase. **(B)** Relative log-expression (RLE) plots of raw expression (top), nSolver-normalized expression with background correction (middle), and nSolver-normalized expression without background correction (bottom) for 90 randomly selected CBCS breast cancer samples, ordered from left to right by increasing per-sample median in the raw expression. The dotted line gives a reference for a deviation of 0.

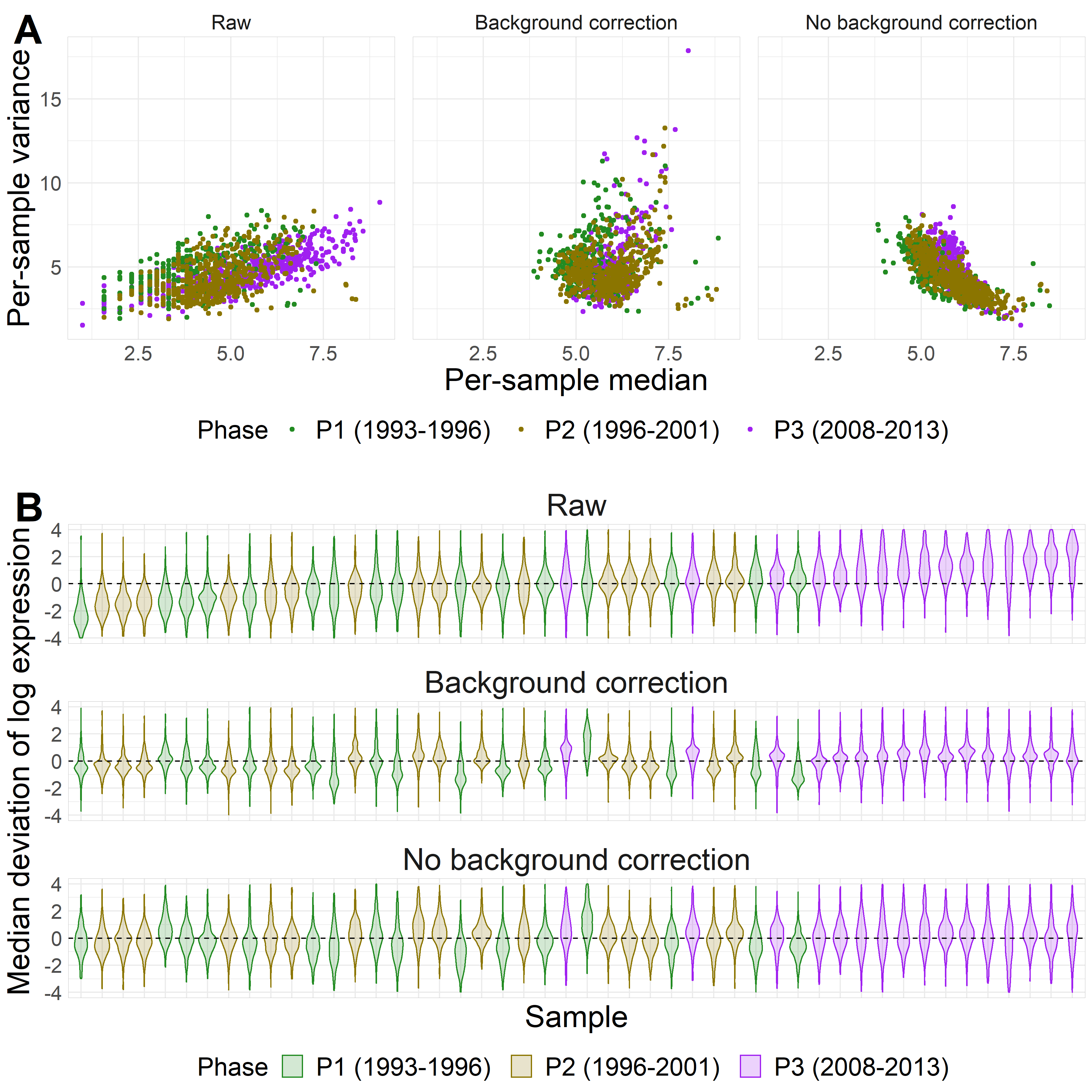

**Figure 2**: Boxplot of percent below the limit of detection (LOD) in endogenous genes ($Y$-axis) over varying numbers below the LOD in the 11 housekeeping genes ($X$-axis), colored by various QC flags

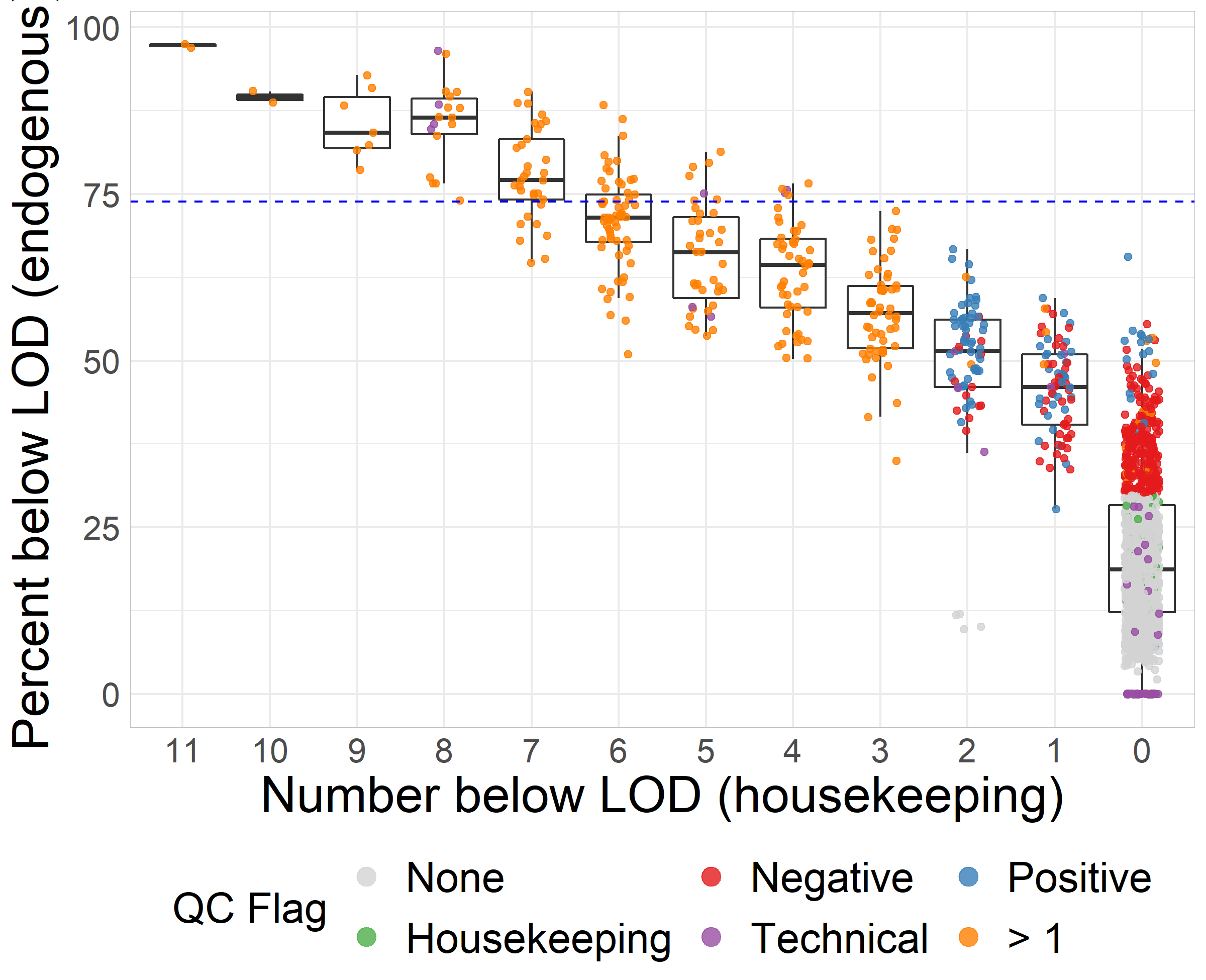

Percent below LOD

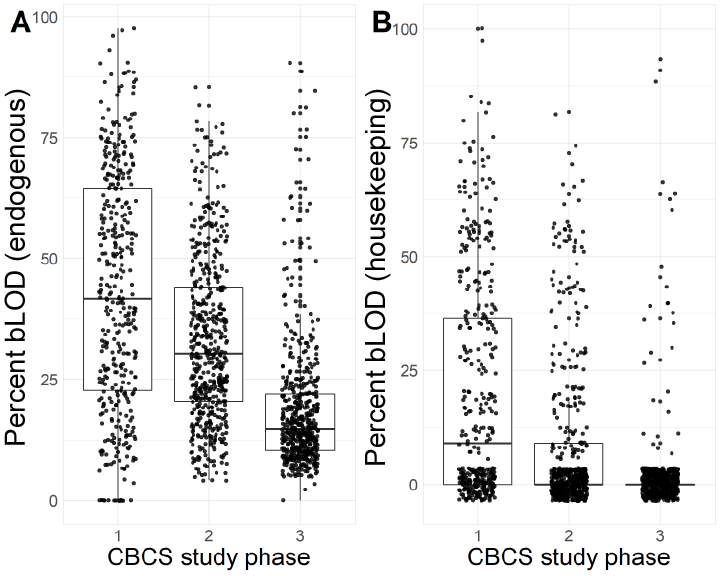

**Figure 3**: Boxplots of percent below the LOD per sample by CBCS study phase with percent bLOD of 406 endogenous genes (A) and percent bLOD of 11 housekeeping genes (B).

Percent below LOD

**Figure 4**: Scatter plots of first two principal components of raw, nSolver-, RUVSeq-, NanoStringDiff-, and RCRnorm-normalized CBCS expression data colored by study phase (A) and PAM50 subtype call (B). PC1 ($X$-axis) captures the maximum variation in expression (approximately 9-12% across all datasets), and PC2 ($Y$-axis) captures the second most (approximately 3-4%).

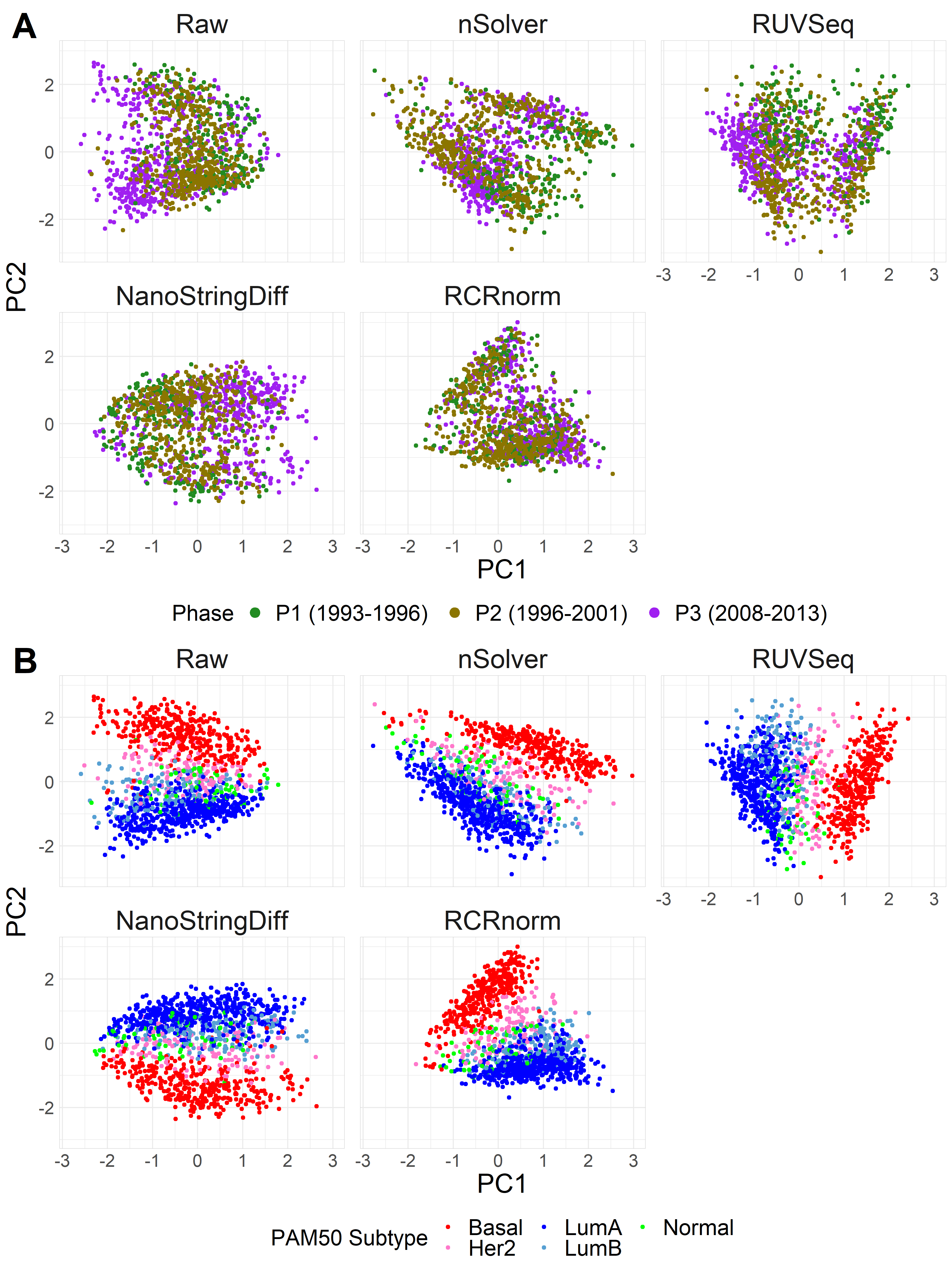

**Figure 5**: Boxplots of silhouette widths of raw, nSolver-, RUVSeq-, NanoStringDiff-, and RCRnorm-normalized CBCS expression data colored by ER status (A) and study phase (B).

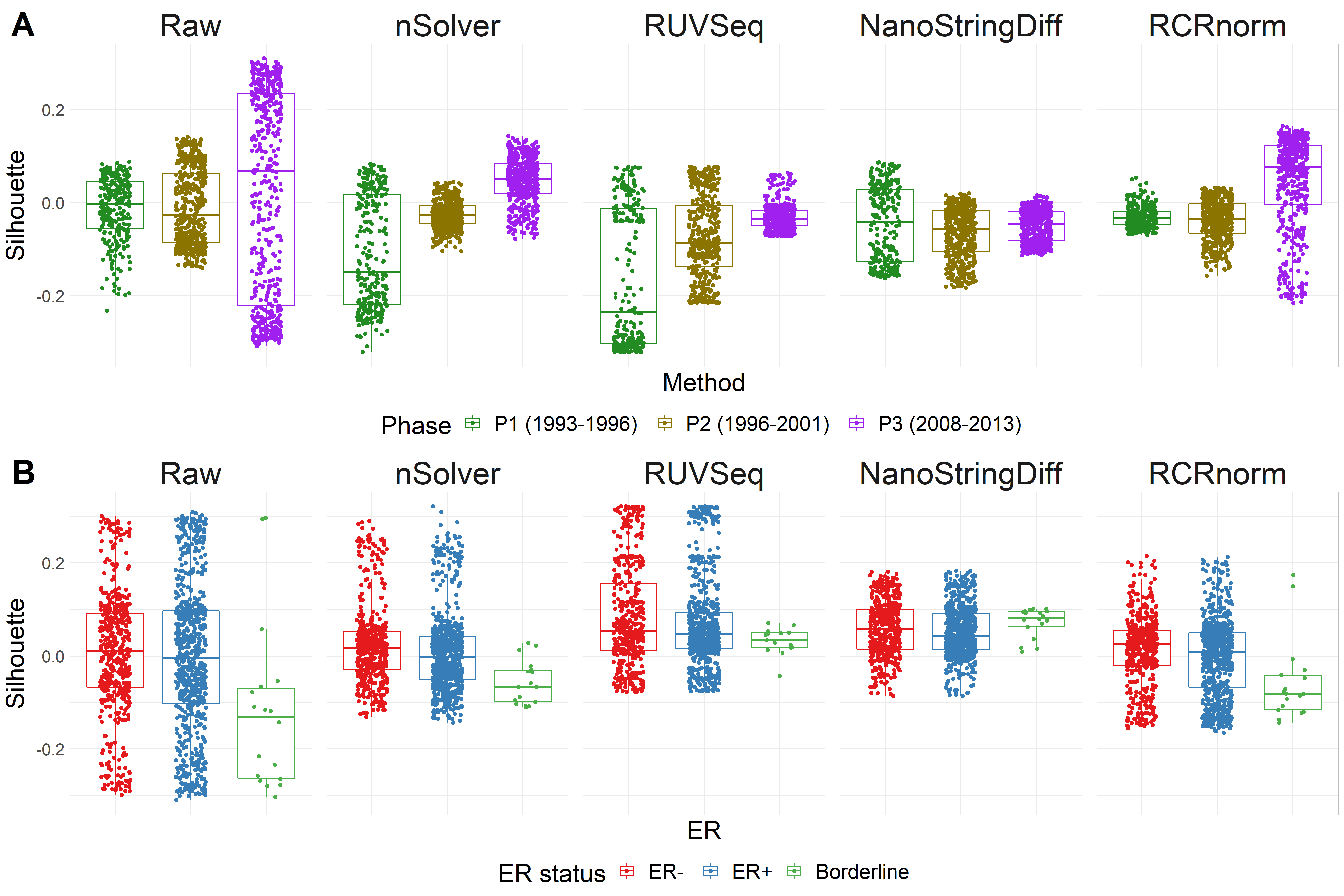

|  |  | **PAM50 subtype calls on nSolver-normalized expression** | | | |  |
| --- | --- | --- | --- | --- | --- | --- |
|  |  | **Basal-like** | **Her2-Enriched** | **Luminal A** | **Luminal B** | **Total** |
| ***PAM50 subtype calls on RUVseq-normalized expression*** | **Basal-like** | 357 | 1 | 1 | 0 | 359 |
|  | **Her2- Enriched** | 0 | 111 | 10 | 0 | 121 |
|  | **Luminal A** | 0 | 12 | 442 | 31 | 485 |
|  | **Luminal B** | 0 | 5 | 38 | 126 | 169 |
|  | **Total** | 357 | 129 | 491 | 157 | 1134 |
|  |  |  |  | **Inter-Rater Agreement (K)=0.87** | | |

**Table 3**: Confusion matrix of PAM50 calls using nSolver-normalized and RUVSeq-normalized expression

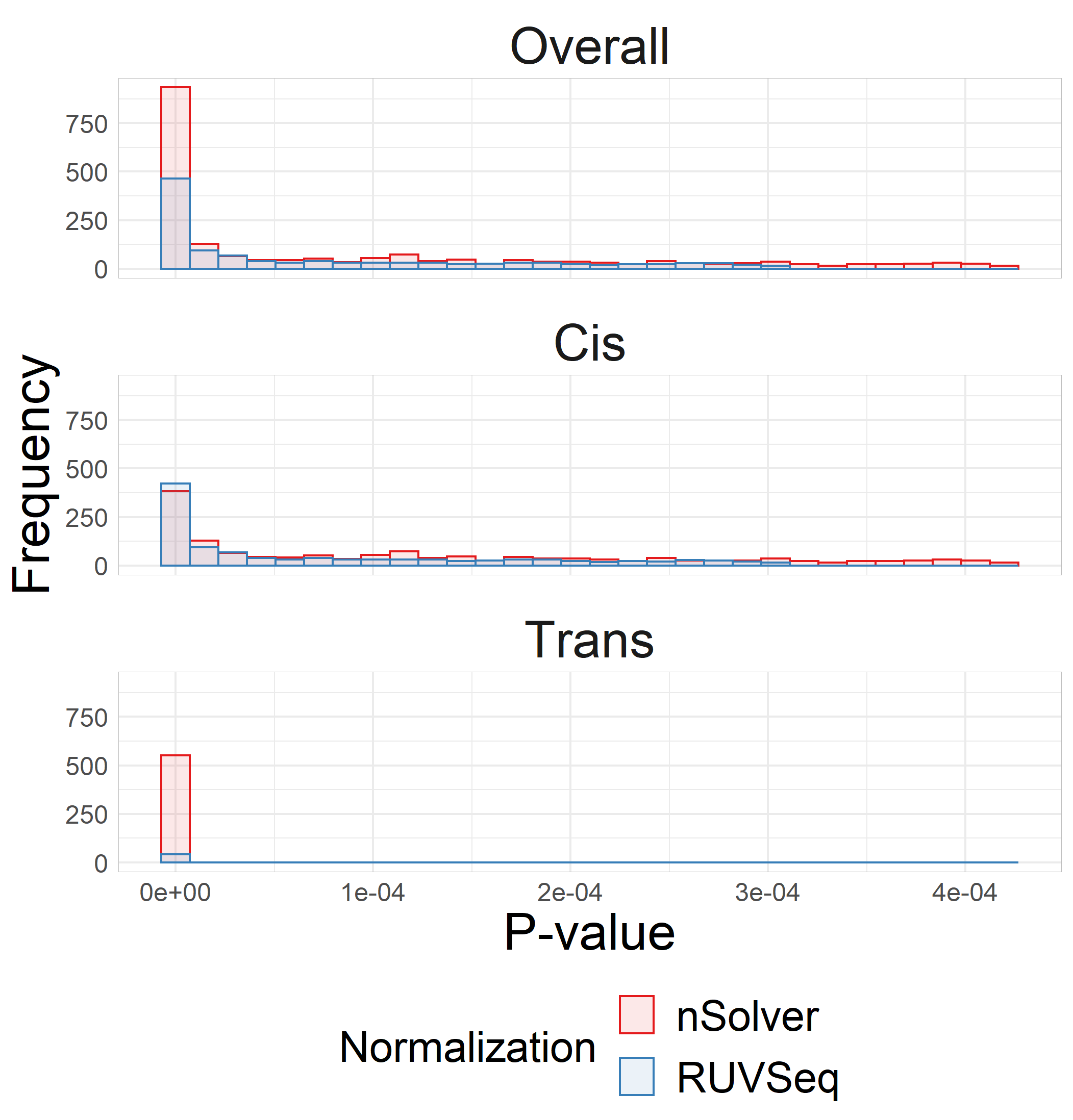

**Figure 6**: Histograms of raw $P$-values of eQTL associations using nSolver-normalized (red) and RUVSeq-normalized (blue) data across overall (top), cis-eQTLs only (middle), and trans-eQTLs only (bottom) for eQTL associations with FDR-adjusted $P<0.05$.

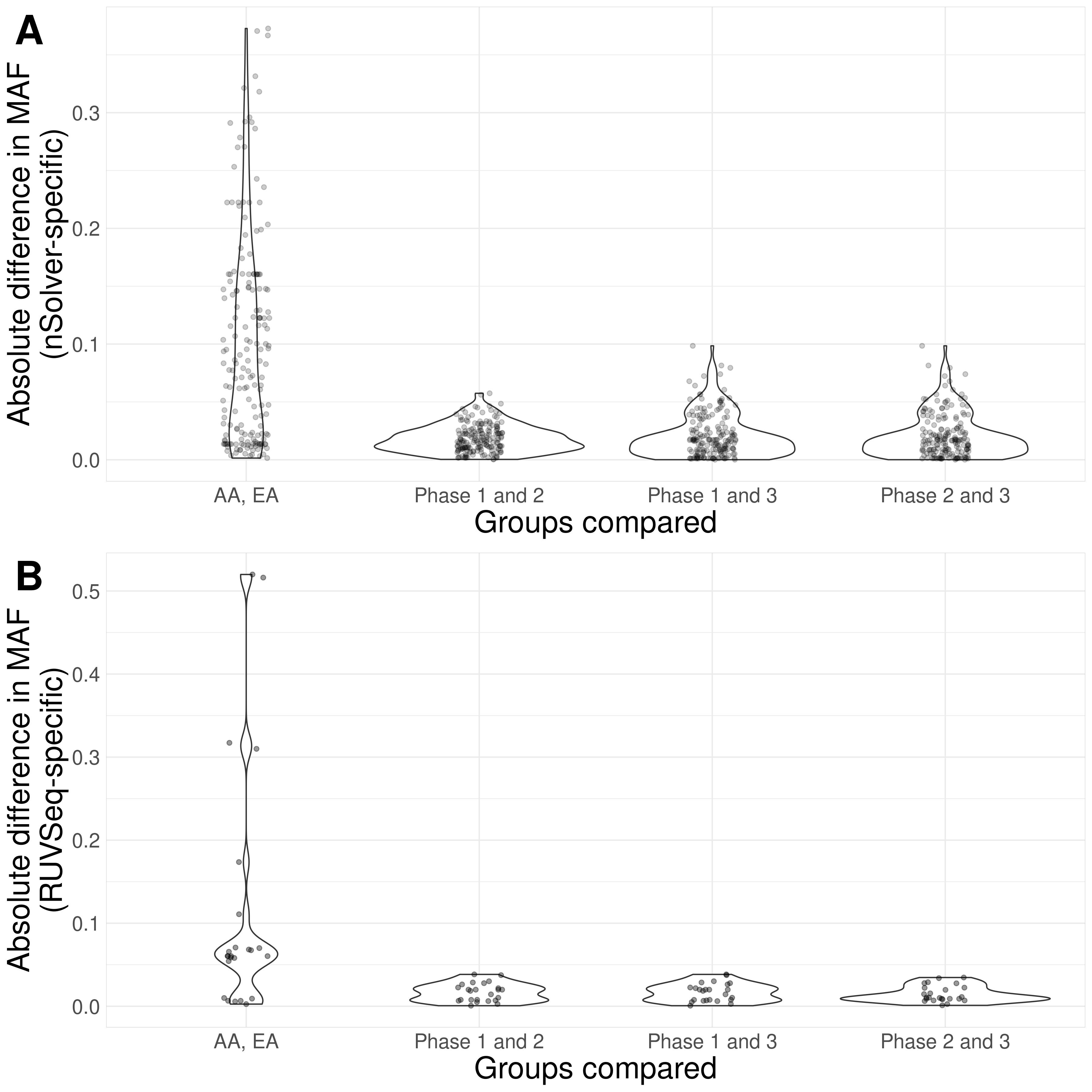

**Figure 7**: Violin plots of absolute differences in minor allele frequencies of trans-eSNPs specific to nSolver-normalized data (A) and RUVSeq-normalized data (B) between groups of African ancestry women (AA) and European ancestry women (EA) and between the three study phases.

**Figure 8**: Directed acyclic graph hypothesizing how spurious trans-eQTLs arise using insufficiently normalized expression. The assumption in the additive linear eQTL model is that the outcome (expression) contains minimal influence from technical variables, like study phase. However, in the case of insufficiently normalized expression that shows variation across study phase, if certain genotypes (trans-SNPs) have differences in minor allele frequency across study phase, study phase serves as an unmeasured confounder. These differences with study phase then bias the association detected between the trans-SNP and expression, causing a spurious association to be detected via eQTL analysis.

trans-SNP

genotypes

Study phase

Insufficiently normalized expression

Spurious trans-eQTL

**Figure 9**: Heatmap of nSolver-normalized (left) and RUVSeq-normalized (right) expression of 417 breast cancer-related genes with hierarchical clustering of samples (horizontal) and genes (vertical). Samples are classified as Basal-like (red), HER2-enriched (pink), luminal A (dark blue), luminal B (light blue), and normal-like (green). The left heatmap uses nSolver-normalized normalized data without quality control based on post-normalization visual inspection. The blue arrow indicates 14 samples without any pre- or post-normalization quality control flags, as outlined in Figure 1, but show deviations from expression patterns.

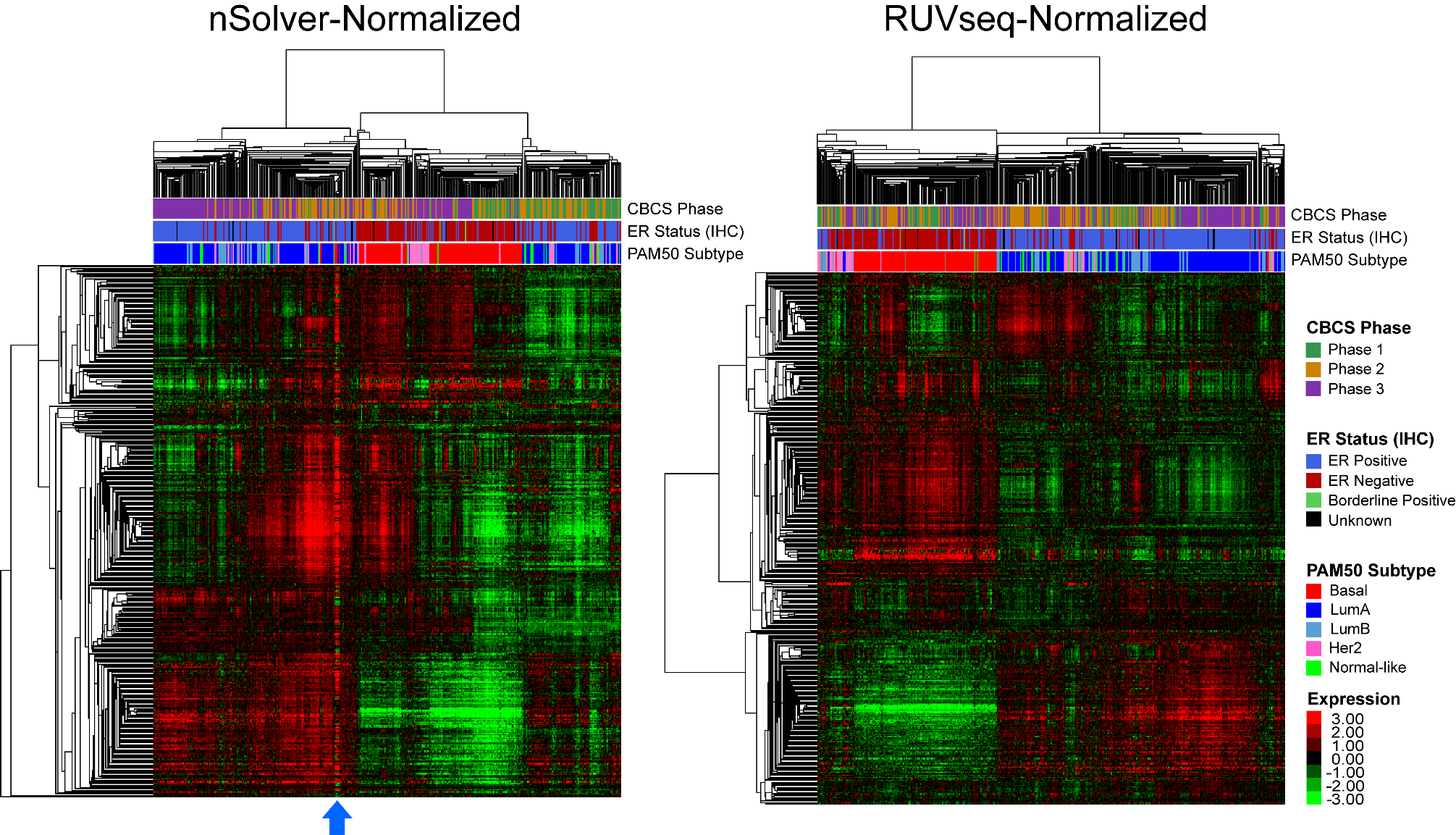

**Figure 10**: Relative log-expression (RLE) plots of raw expression (A), nSolver-normalized expression (B), and RUVSeq-normalized expression (C) for Sabry et al’s natural killer Nanostring expression profile. Boxplots are colored by various treatment groups.

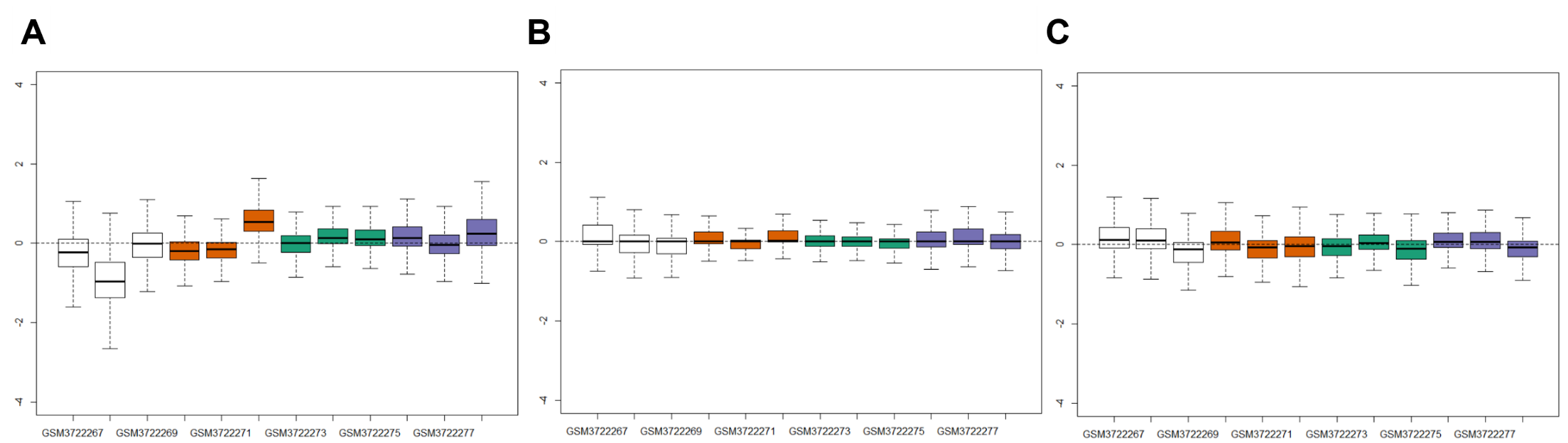

**B**

**Figure 10**: Scatter plots of first two principal components of nSolver-normalized expression data (left) and RUVSeq-normalized data (right) colored by (A) low or high DV300 value and (B) tumor stage for kidney tumor data.
